# An Integrated Neuroimmune Assembloid Model to Advance Neurodegenerative Disease Studies

**DOI:** 10.64898/2026.04.28.721373

**Authors:** Iratxe Ruiz-Formoso, Ikerne Martin-Ferrer, Nerea Urrestizala-Arenaza, Estibaliz Capetillo-Zarate, Fabio Cavaliere, Paula Ramos-Gonzalez

## Abstract

Brain organoids are three-dimensional cultures derived from human pluripotent or embryonic stem cells that recapitulate key genetic, biochemical, and molecular features of the human brain. They provide a powerful platform for studying human brain development and modeling genetic neurological disorders. However, their application to age-dependent neurodegenerative diseases remains limited, largely due to the absence of standardized methods for incorporating functional microglia, critical regulators of neuroinflammation and disease progression. Here, we describe a strategy for generating neuroimmune assembloids, brain organoids containing functional glial cells capable of mounting inflammatory responses. By introducing hematopoietic progenitor cells into developing brain organoids, we enable their *in situ* maturation into microglia-like cells that persist in culture for up to one month. These cells exhibit hallmark microglial behaviors, including morphological remodeling, migration, phagocytosis and transcriptional changes in response to inflammatory stimuli. Together, these immunocompetent-like brain organoids provide a promising and versatile platform for investigating neuroimmune interactions and neuroinflammatory mechanisms underlying age-related neurodegenerative diseases.

## Introduction

Microglia, the resident immune cells of the central nervous system (CNS), originate from macrophage progenitors generated in the yolk sac that migrate into the developing brain around the fourth gestational week [1,2] During brain development, microglia are essential for establishing and maintaining neuronal function, as they support neurogenesis, regulate programmed cell death, and mediate synaptic pruning, thereby contributing to tissue homeostasis. In addition, microglia play a central role in neuroinflammatory processes associated with neurodegenerative diseases [3].

Guided brain organoids are designed to recapitulate specific embryonic developmental trajectories and are composed exclusively of neuroectoderm-derived cells. The differentiation of mesoderm-derived lineages, including microglia, is actively suppressed through dual SMAD inhibition of bone morphogenetic protein (BMP) and transforming growth factor beta (TGF-β) signaling pathways. Consequently, generating cerebral organoids containing physiologically relevant populations of mature microglia has remained a major challenge in the field [4]. Recent approaches using mild dual SMAD inhibition, achieved by reducing concentrations of the bone morphogen protein 4 (BMP4) inhibitor dorsomorphin, have enabled the simultaneous differentiation of ectodermal and mesodermal lineages, allowing endogenous microglia-like cells to emerge [5]. However, this strategy introduces substantial variability and heterogeneity that is highly sensitive to dorsomorphin dosage. Alternative methods rely on gene induction systems to overexpress PU.1, a pan-macrophage transcription factor, which can promote microglia-like differentiation even under neuroectoderm-favoring conditions, albeit with inconsistent efficiency. The most commonly used approaches instead involve the exogenous addition of microglia to developing human cerebral organoids, either from postmortem human tissue [6] or immortalized human microglial cell lines [7]. While these methods yield organoids containing cells that express canonical microglial markers, the resulting microglia often exhibit limited parenchymal invasion, diminished or hyper-neuroinflammatory responsiveness, and lack a homeostatic transcriptional profile. Despite numerous efforts to incorporate microglia into guided brain organoids [4,8], the robust generation of immunocompetent microglia within brain organoids remains an unmet challenge. Immunocompetence refers to a functionally mature microglial state characterized by regulated inflammatory responses, efficient phagocytic activity, dynamic transitions between branched and amoeboid morphologies, mosaic-like spatial distribution, and homeostatic gene expression signature.

Here, we present a strategy to generate neuroimmune assembloids by introducing primed induced pluripotent stem cell-derived hematopoietic progenitor cells (iHPCs) into developing brain organoids. Within one month, iHPCs differentiate in situ into microglia that robustly infiltrate the organoid tissue and display diverse morphologies. These microglia respond functionally to neuroinflammatory stimuli and support the long-term viability and functionality of brain organoid cultures.

## Materials and Methods

### iPSC-derived cerebral organoids (hCOs)

hCOs were generated from iPSC-derived embryoid bodies (EBs) using a slightly modified protocol based on the commercial serum-free StemDiff Cerebral Organoid kit (STEMCELL, Grenoble, France ref. 08570, Fig. 1A). Briefly, EBs were generated by seeding 4,000 iPSCs from the standardized KOLF2.1J line (Jackson Laboratory, USA) in AggreWell plates (STEMCELL Technologies). For this initial step, the EB Formation Medium of the kit was replaced with SMADi Neural Induction Medium (STEMCELL ref. 08581). On the seeding day, ROCK inhibitor Y-27632 (Merck, Madrid, Spain) was added to the medium. Medium changes were performed daily for the following 5 days. During this phase, precursor cells in the EBs were directed toward a neuroepithelial fate using dual SMAD inhibition to suppress non-neural differentiation.

**Figure 1.**
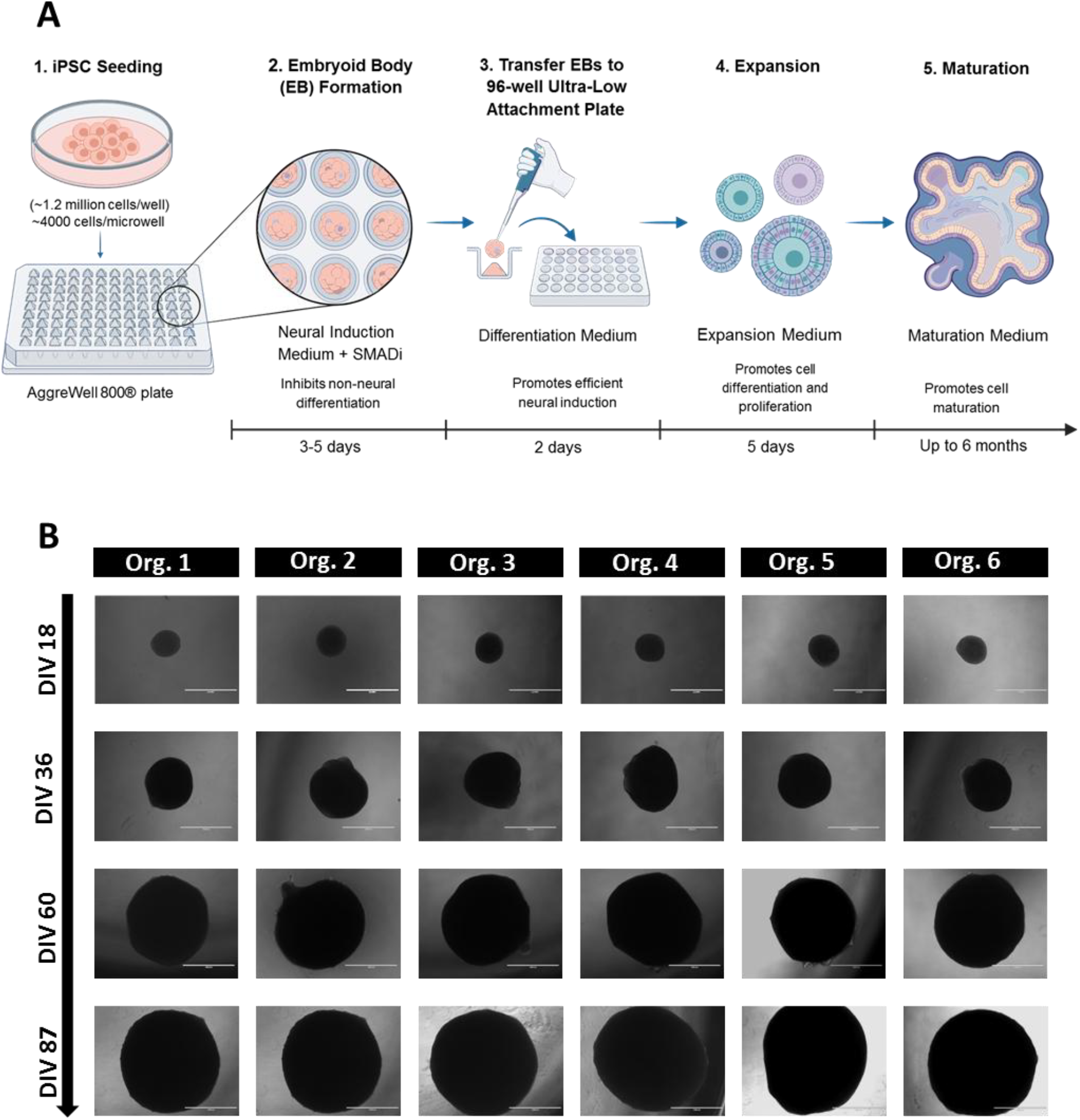
Protocol for generating cerebral organoids from the KOLF2.1J iPSC line. A) Schematic of brain organoid generation starting from iPSC seeding. Detailed protocol is provided in the Materials and methods section. B) Brightfield images of 6 different organoids cultured for 87 days. The homogeneity in size and morphology indicates methodological reproducibility. Scale bar=800 μm.

After 5 days, each EB was transferred into a 96-well round-bottom ultra-low attachment plate (Corning, NY, USA; ref. 7007) and from this point onward, the StemDiff Cerebral Organoid Kit protocol was followed according to the manufacturer’s instructions: EBs were cultured in differentiation medium for 2 days, on day 7, organoids were transferred to Expansion Medium, and on day 10 they were switched to Maturation Medium and placed on an orbital shaker (Infors HT, Switzerland) at 57 rpm, 37°C, and 5% CO_2_.

Neural precursor cell identity was validated by immunofluorescence for PAX6, SOX2, and Vimentin and maturation was assessed by immunofluorescence for neuronal markers (βIII-tubulin, MAP2, synaptophysin), astrocytic markers (GFAP, AQP4), and oligodendrocytic markers (MBP, SOX10). Functional maturation was further evaluated by spontaneous electrical activity using high-density multi-electrode arrays (HD-MEA). Up to 150 organoids were generated from a single iPSC line. Quality control was performed by immunofluorescence and RT-PCR in a subset of 15 organoids (Table 1).

**Tab.1.**
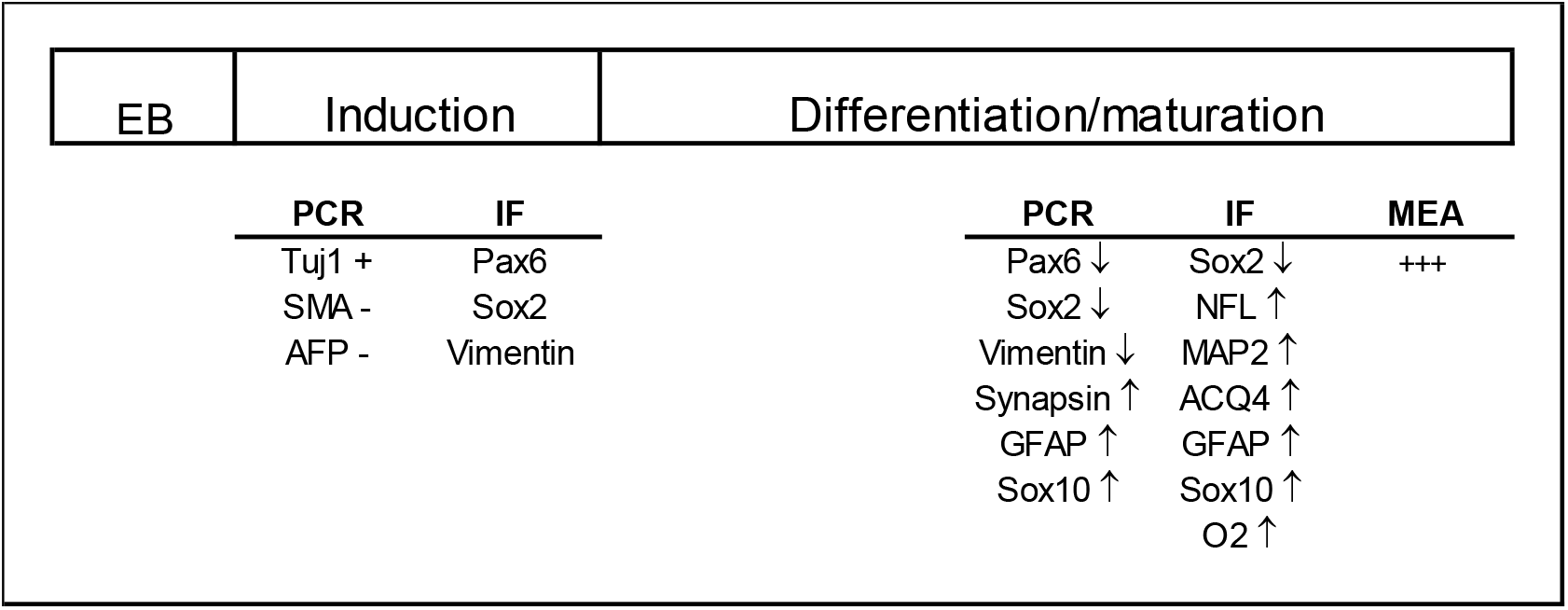
QC of organoid culture is performed by immunofluorescence (IF), PCR and MEA at the end of induction and maturation (Tuj1, ectoderm; SMA, mesoderm; AFP, endoderm): embryoid bodies (EB) are not characterized.

### iPSC-derived microglia (iHPC) 2D model

Microglia were generated from iPSCs using a commercial STEMCELL Technologies protocol (Fig. 3A). A three-stage differentiation system was used to generate hematopoietic progenitor cells (iHPCs) (STEMCELL, STEMdiff™ Hematopoietic Kit cat. 05310), microglial precursors (STEMCELL, STEMdiff™ microglia differentiation Kit cat. 100-0019), and mature microglia (STEMCELL, STEMdiff™ microglia maturation kit cat. 100-0020).

During the first 3 days, cells were cultured in STEMdiff™ Hematopoietic Supplement A to induce mesoderm specification. During the following 9 days, STEMdiff™ Hematopoietic Supplement B was used to promote differentiation into hematopoietic progenitors. After 12 days, floating iHPCs were collected and seeded onto Matrigel-coated 6-well plates (Corning, NY, USA) and differentiated into microglia using STEMdiff™ Microglia Differentiation Kit for 24 days and 6-10 days in STEMdiff™ microglia maturation kit.

### Neuroimmune assembloids

To generate neuroimmune assembloids, iHPCs were primed for 2 days in microglia differentiation medium (STEMCELL Technologies) before assembling in hCOs. Subsequently, 20,000/30,000 iHPCs were added to 30–40 day-old hCOs by gentle centrifugation (500 × g for 3 min) and matured within organoids on a Celltron orbital shaker at 57 rpm, 37°C, and 5% CO_2_. Original organoid medium was replaced under gradual medium transitions performed every 2 days (30%/70%, 50%/50%, 70%/30%, and 100% ratios of organoid/microglia media). Assembloids were further maintained for up to 30 days (Fig. 4A).

### Phagocytosis assay

Phagocytosis assays were performed as previously described (Beccari et al. [9]). SH-SY5Y VAMPIRE cells (provided by Dr. A. Sierra) were labelled with tFP602 red fluorophore (InnoProt, P20303) and treated with staurosporine (3 μM, 4 h; Sigma-Aldrich, S4400) to induce apoptosis and cell fragmentation. Phagocytosis was assessed by incubating microglia with apoptotic fluorescent debris for 3 h, followed by fixation to evaluate engulfment of labelled cells.

### Immunofluorescence

Organoids were fixed overnight at 4°C in 4% paraformaldehyde (PFA; Merck, Madrid, Spain), washed in PBS, embedded in 4% agarose, and sectioned at 70 µm using a vibratome. Free-floating sections were permeabilized in PBS containing 0.5% Triton X-100 for 15 min and blocked in 5% goat or donkey serum in PBS with 0.1% Triton X-100. Sections were incubated overnight at 4°C with primary antibodies (Table 2). After washing, samples were incubated with Alexa Fluor–conjugated secondary antibodies for 2 h at room temperature. Sections were counterstained with DAPI and mounted using Fluoromount (Sigma-Aldrich, Madrid, Spain). Imaging was performed using a Leica Stellaris confocal microscope at the Achucarro Basque Center for Neuroscience Imaging Facility.

**Tab. 2.**
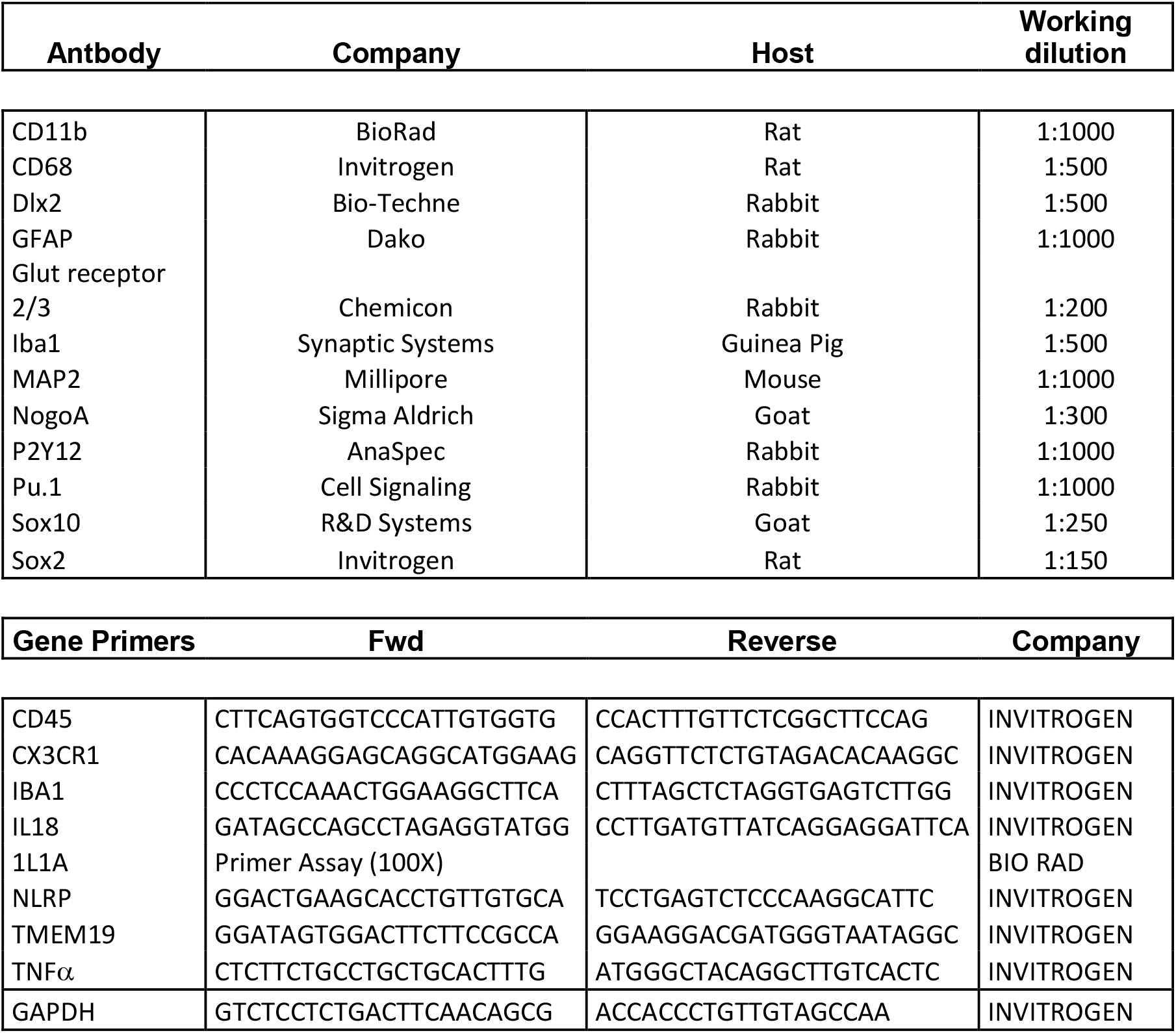
Antibodies and primers used for 2D microglia, brain organoids and neuroimmune assembloids.

### Morphology assay and microglia invasion

The total of five organoids for each condition were analyzed using the open source Software Fiji (Schindelin, J. et al., 2012). Five slices from each organoids immunostained with Iba1 antibody were analyzed with different plug-ins. All measurements were exported for statistical analysis with GraphPad 8.0.

#### Circularity

Images were converted to 8-bit and thresholded to segment individual cells, generating binary masks; the ‘Watershed’ tool was applied when needed to separate touching cells. Shape descriptor “circularity” was calculated as 4π·Area/Perimeter2.

#### CD68 area occupancy

For each segmented cell, thresholded binary masks were generated for the Iba1 and CD68 channels. In this case, shape descriptor “area” was measured. The CD68-positive area was then normalized to the corresponding Iba1-positive area, yielding a CD68/Iba1 area ratio for each cell.

#### Spatial localization

Analysis was performed at different days (from 1 to 30 days) after microglia assembly. Spatial localization was calculated as the percentage of area occupied by labelled microglia vs total area of the organoid, using the Software Fiji. When spatial localization per areas was analyzed, organoid area was divided in three different arbitrary zones: the external, intermediate and central.

### Total RNA and qRT-PCR

Total RNA from 5 organoids was extracted using the commercial kit NZY Total RNA isolation kit (NZYtech, Lisbon, Portugal). The quality of extracted RNA was checked on agarose gel and 1 µg was retro-transcribed at 60°C for 60 minutes using Superscript SSIII (Invitrogen, Madrid, Spain). A 2µl aliquot (about 100ng) of each cDNA was used for real-time quantitative PCR in the BioRad CFX96 thermocycler using the primers listed in Tab.2. ΔCt for each transcript was calculated after normalization with GAPDH transcript.

#### Cytofluorimetry assay

Brain organoids were dissociated at single cell level using papain before cytofluorimetry analysis. Briefly, five organoids were plotted together and triturated in freshly prepared dissociation solution (30 units/mL papain, 125 units/mL DNase I) by pipetting 10 times at room temperature. Cell suspension was incubated at 37°C for 10 minutes on an orbital shaker set to 90-120 rpm and triturated 10 times at room temperature to obtain a suspension consisting primarily of single cells. The entire cell suspension was centrifuged in tubes containing 2 mL of 10 mg/mL ovomucoid protease inhibitor solution and centrifuged at 300 x g for 5 minutes. The cell pellet was resuspended in 500 µL of PBS and filtered through a BD 50 µL sterile cup Filcon (Cat# 340630). 1x10^6^ cells were labelled with 10 µL of Pharmingen™ Propidium Iodide Staining Solution (Cat# 556463) in 500 µL of PBS and analyzed as FSC vs 585/29-PE/DSRed at the Cell Analytics Service of Achucarro Basque Center for Neuroscience using a BD FACS Jazz system (BD Biosciences).

#### Electrophysiology recording

Extracellular electrophysiology was performed by high density multi electrode array (HD-MEA) using the Biocam Duplex (3Brain, Switzerland). One single organoid or assembloid was previously cultured in BrainPhys™ (STEMCELL) for 7 days and seeded in the same medium on a Core Plate 1W-2D chip. Plates were equilibrated for 10 minutes and basal activity recorded for 20 minutes. Electrodes were recorded with High pass filter at 10Hz and spike detection was performed using the PTSD algorithm (peak life time, 2 mS, refractory period, 1 mS). Raw data were analyzed with the Brain 6 Software and electrical activity measured as mean firing rate (spike/second) or number of spikes, whereas synchronicity was evaluated as a connectivity map.

## Results

### Generation and characterization of cerebral organoids

To generate reproducible cerebral organoids with low heterogeneity, we employed a standardized commercial protocol (see Materials and Methods) using the reference human iPSC line KOLF2.1J (Jackson Laboratory). Human cerebral organoids (hCOs) were generated using a three-stage differentiation protocol adapted from Lancaster et al. [11] (Fig. 1A). Embryoid bodies (EB) were formed in standardized microwells (800 µm diameter) to promote uniform neuroepithelial organization and subsequently differentiated and matured into hCOs for up to 240 days in culture. As shown in Fig. 1B, we obtained a morphologically homogeneous organoid culture. Within 20 days, hCOs displayed characteristic neuroepithelial rosette structures expressing the ventricular zone markers SOX2, indicative of proliferative neural progenitor regions (Fig. 2A). Neuronal differentiation was evident from day 30 onward, as demonstrated by the expression of MAP2 in dendrites and SMI31 in phosphorylated neurofilaments (NFL) throughout the differentiation timeline. In contrast, expression of the AMPA glutamate receptor Glu2/3 and the

**Figure 2.**
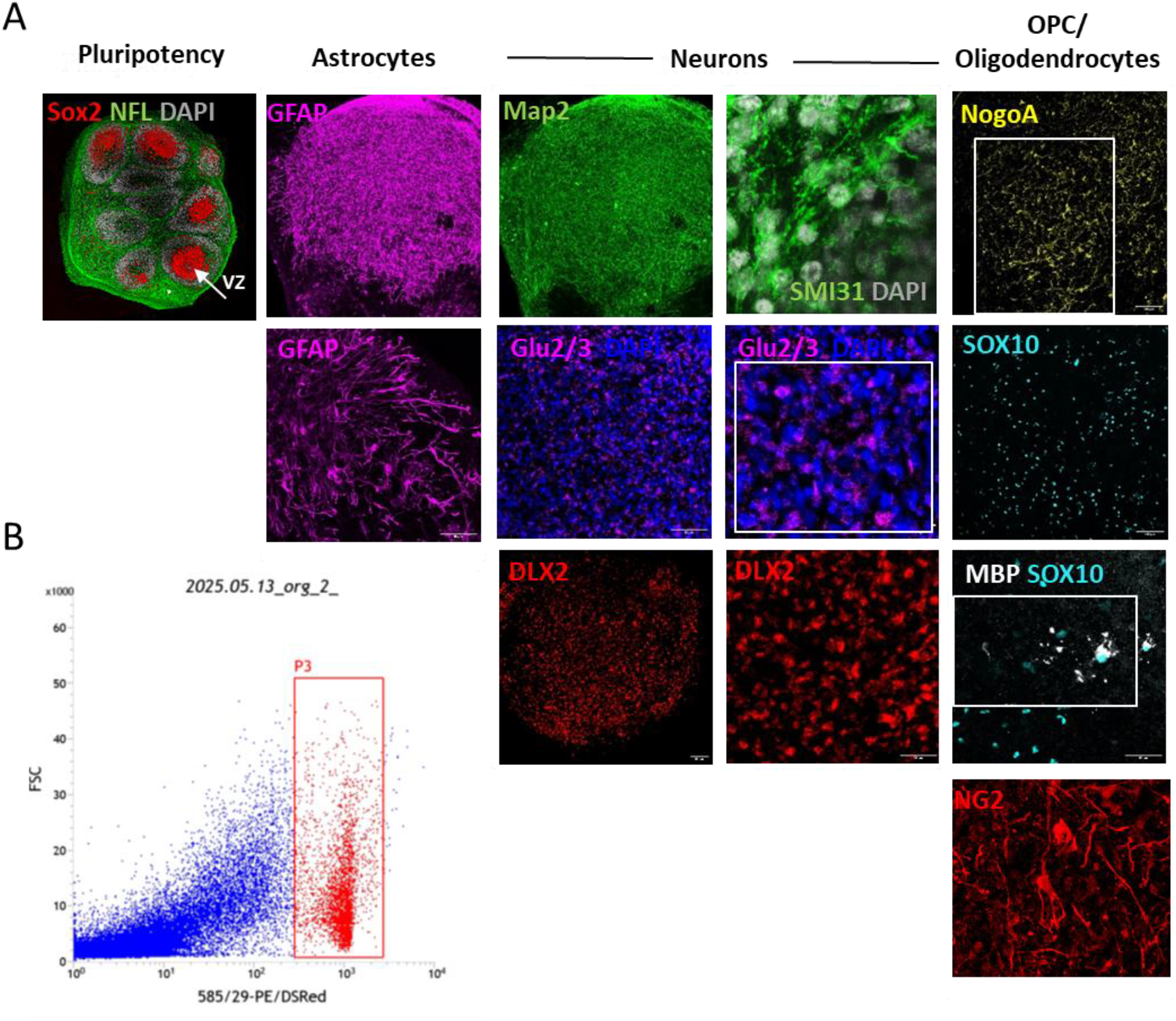
Characterization of human cerebral organoids. A) hCOs were fixed in 4% paraformaldehyde after different days *in vitro* (DIV) and analyzed by immunofluorescence using markers for different cell types. Sox2, a pluripotency marker, was stained after 15 DIV, arrow indicates the ventricular zone (VZ); GFAP, astrocyte marker, was detected after 40 DIV and visualized with two different magnifications (10x and 40x); the general markers for staining mature neurons, MAP2 and neurofilament SMI31 (NFL), were respectively visualized with 10x and 40x objectives after 30 DIV, whereas Glu2/3 (20x and 40x magnification) and Dlx2 (20x and 40x magnification) were stained after 40 DIV. Markers for oligodendrocyte precursor cells (Sox10, NogoA and NG2) and mature oligodendrocytes (MBP) were visualized after 75 DIV. Squares in Glu2/3, NogoA and MBP represent the zoom-in of the images. B) Single-cell analysis by cytofluorimetry assay to quantify cell death after propidium iodide staining. Death cells are gated as P3.

GABAergic lineage marker DLX2 was detected only after 40–60 days in culture, reflecting progressive neuronal maturation. Astrocytic differentiation, assessed by GFAP expression, was observed after day 40. Although markers of oligodendrocyte lineage cells, including NogoA, SOX10, MBP and NG2 indicated the presence of both immature and mature oligodendrocytes, myelination remained sparse (Fig. 2A). Finally, overall cell viability in mature hCOs was assessed by dissociating organoids into single cells and quantifying cell death using propidium iodide staining followed by flow cytometry analysis. This analysis revealed low levels of cell death, ranging from 10–18%, confirming the robustness and viability of the organoid cultures (Fig. 2B).

### Generation of human iPSC-derived microglia and neuroimmune assembloids

Before assembling human microglia into hCOs, we first assessed their immunocompetence in a two-dimensional (2D) culture system. To generate immunocompetent microglia, defined by regulated inflammatory signaling, phagocytic activity, and reversible transitions between ramified and amoeboid morphologies, we differentiated the human iPSC line KOLF2.1J into adult microglia. The differentiation protocol (Fig. 3A) supports mesodermal specification, subsequent generation of iHPCs, and full maturation into adult microglia within 30-40 days. Mature microglia cultured in 2D expressed canonical microglial markers, including P2Y12, a homeostatic marker characteristic of resting microglia, as well as the activation markers IBA1 and CD11b, and the myeloid transcription factor PU.1 (Fig. 3B). Microglial functionality was confirmed using a phagocytosis assay [9]. SH-SY5Y neuroblastoma cells were pre-labeled with the red fluorophore tFP602 and induced to undergo apoptosis with 3mM staurosporine. Fragmented apoptotic cells (red labelled) were then incubated with mature microglia, and phagocytic uptake was visualized by red fluorescence within microglial cells three hours after exposure (Fig. 3C).

**Figure 3.**
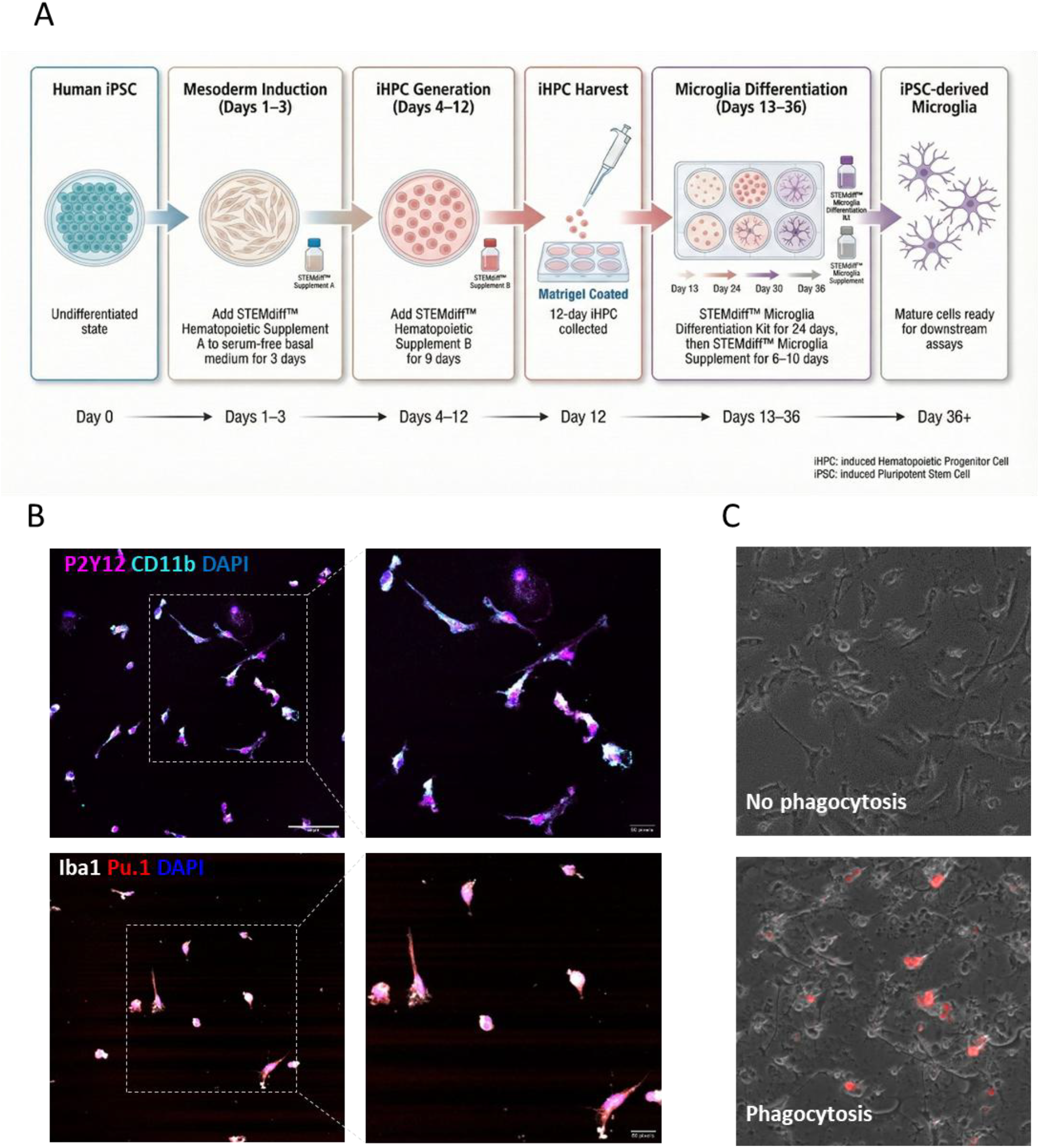
2D microglia protocol (A) and characterization (B-C). B) Microglia differentiated and matured after 35 days were stained for the expression of homeostatic (P2Y12 in purple) and activated genes (CD11b, cyan; Iba1, white) and the transcription factor PU.1 (red). Scale bar=50μm. Images on the left are zoomed in the right panels. C) Phagocytosis analysis using Vampire cells. Red fluorescence inside the microglia indicates phagocytosis.

During human development, HPCs begin migrating from the yolk sac to the developing brain around the fifth post-conceptional week and continue to migrate within the parenchyma of different brain regions until infancy and early childhood [1]. During their expansion, microglia differentiate into distinct specialized populations depending on the brain region. This differentiation and specialization are further supported by astrocytes, which provide essential trophic factors such as cholesterol, CSF-1, TGF-β, and IL-34 (as discussed below). Based on these developmental principles, to generate functional neuroimmune assembloids we differentiated hematopoietic precursor cells within 60-day-old brain organoids when mature astrocytes are already present (Fig. 4A). Prior to incorporation into brain organoids, between 20,000 and 30,000 iHPCs were primed in microglial differentiation medium for 2 days (see detailed protocol in Supplementary Methods). Primed iHPCs were then centrifuged together with a single organoid at 300×g for 5 minutes, and microglial invasion, differentiation, and functionality were monitored within the organoids over a period of 30 days. To promote microglial survival during the early phases of differentiation and maturation into the organoid, we gradually changed the culture medium, starting with 100% microglia medium and replacing 25% with organoid medium every 2 days. (Fig. 4A). Microglial invasion was assessed by immunofluorescence using an Iba1 antibody at 1, 4, 7 and 14 days after assembly. We observed a progressive migration of microglia displaying diverse morphologies, ranging from branched to rounded/amoeboid forms (Fig. 4B-C), moving from the periphery toward the core of the organoids. The area occupied by microglia reached a maximum 7 days after assembly, followed by a slight decrease at 14 days (Fig. 5A-B) and hyperactivated rounded morphology at 30 days (data not shown). To further characterize microglial invasion, we quantified the area occupied by microglia over 14 days within three arbitrarily defined zones: external, intermediate, and central (Fig. 5C-D). One day after assembly (1 DIV), microglia were almost exclusively localized in the external zone, with only residual cells present in the intermediate zone. Four days after inclusion, microglia had migrated inward, with approximately 58% remaining in the external zone, 31% in the intermediate zone, and 11% reaching the central zone. By 14 DIV, we observed a continued redistribution toward the organoid core, with microglia distributed approximately 38% in the external zone, 42% in the intermediate zone, and 20% in the central zone. The inclusion of microglia in neuroimmune assembloids also had functional consequences for overall brain organoid maturation. Fourteen days after microglial incorporation, high-density multi-electrode array (HD-MEA) analysis revealed a more functional basal electrical activity in neuroimmune assembloids compared with organoids without microglia (Fig.6). The activity map observed during HD-MEA registration (Fig. 6A) showed a functional increase of the mean firing rate (Fig.6B) as well as higher number of spikes (Fig. 6C). Overall, these features generated a more complex connectivity map in neuroimmune assembloids compared with organoids lacking microglia (Fig. 6D), suggesting enhanced synchronization and neuronal network functionality.

**Figure 4.**
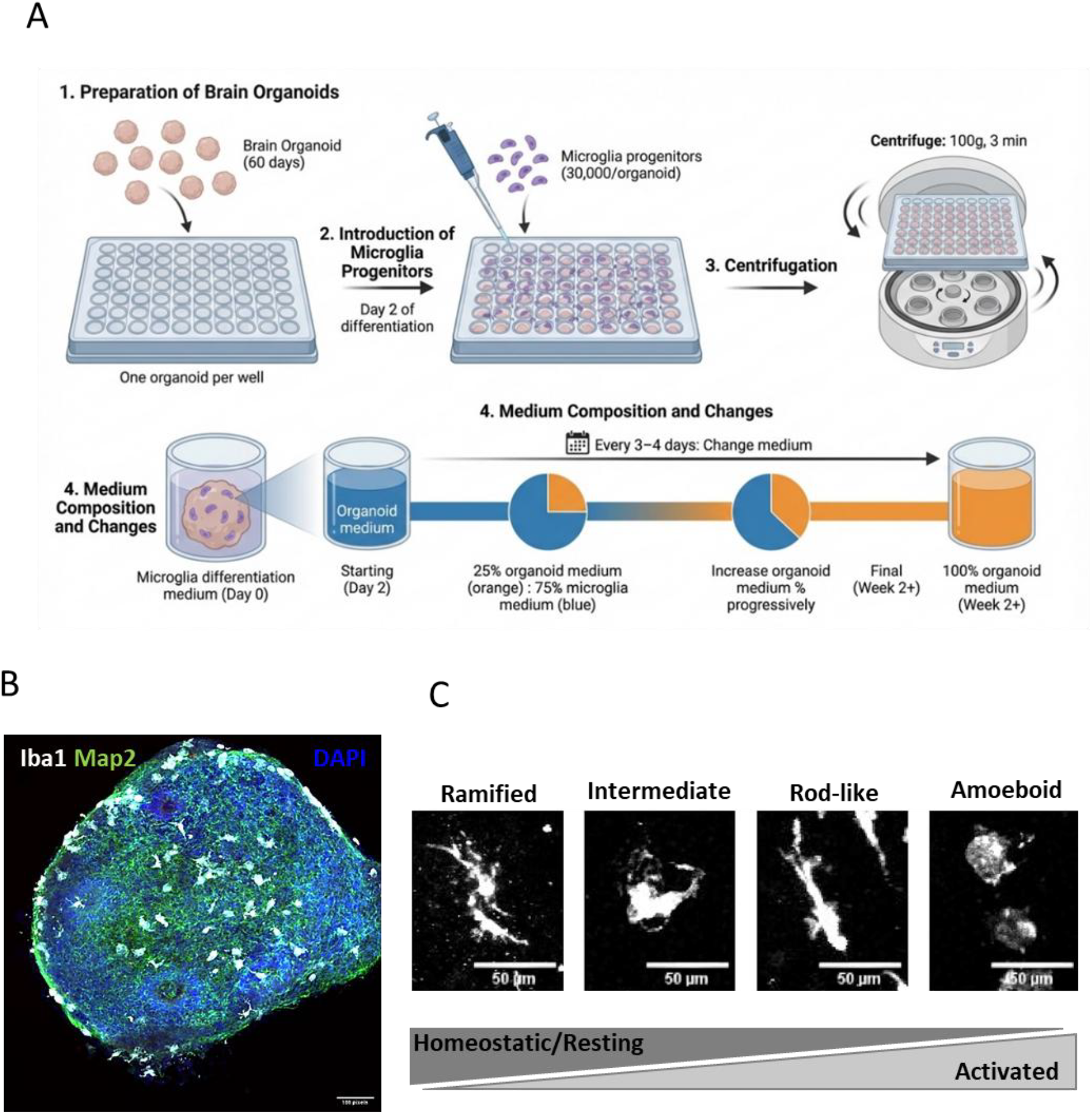
Neuroimmune assembloids protocol (A). B) Representation of a 75 DIV hCO 14 days after microglia assembly. Microglia are stained in white using a Iba1 antibody. The neuronal marker MAP2 for neuronal dendrites is shown in green, and nuclei are counterstained with DAPI in blue. C) Neuroimmune assembloids show Iba1-positive microglia with distinct morphologies, suggesting the presence of both resting and activated microglia.

**Figure 5.**
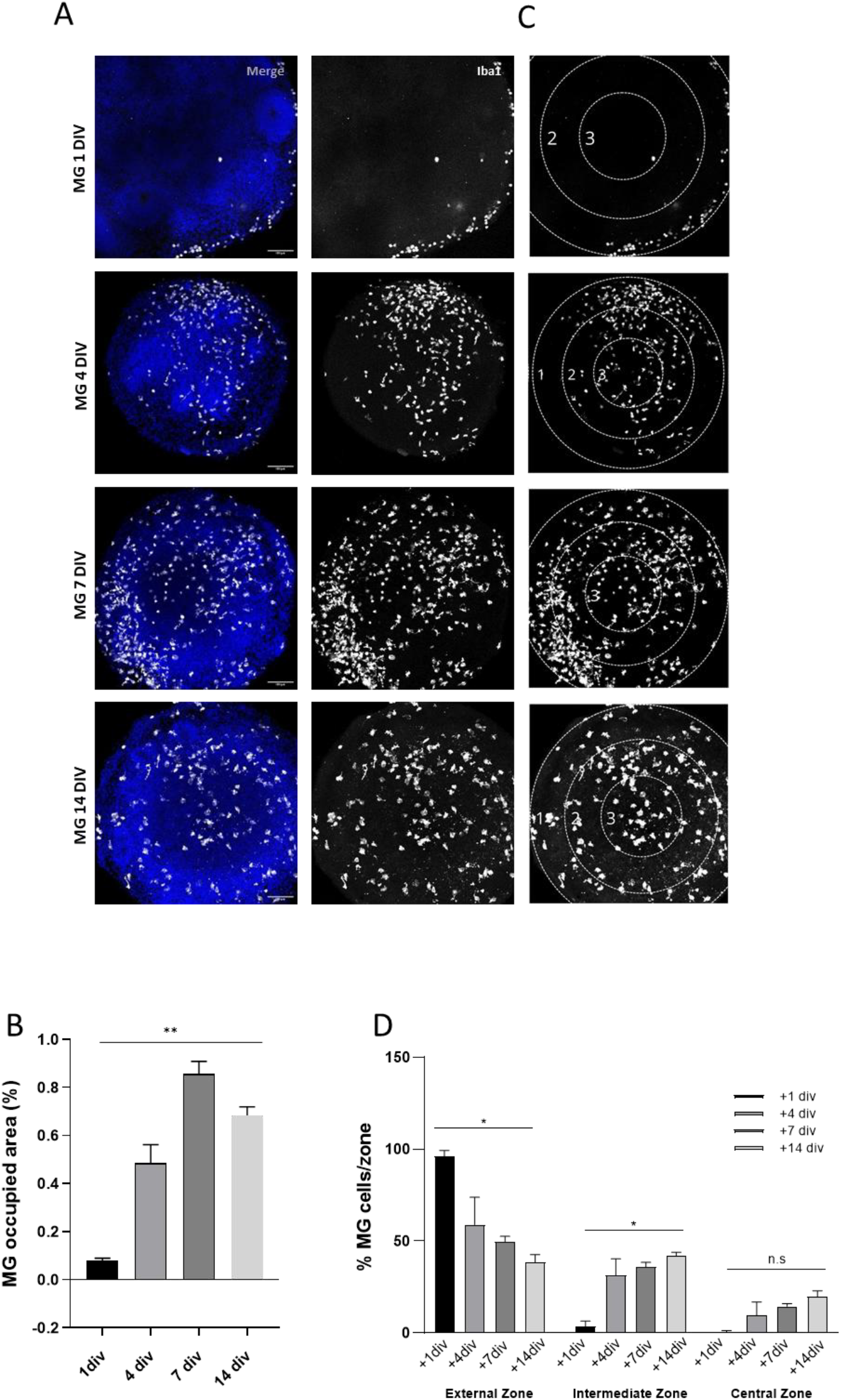
Microglia invasion in neuroimmune assembloids. A) Neuroimmune assembloids were fixed 1, 4, 7 and 14 days after microglia (MG) assembly in hCO and stained by immunofluorescence for Iba1. (B) Quantification of area occupied by microglia (%) different time points after assembly. (C) Number of microglia in different zones of neuroimmune assembloids (1, external; 2, intermediate; 3, central zone). D) Quantification of microglia invasion across area 1, 4, 7 and 14 days (DIV) post-assembly.

**Figure 6.**
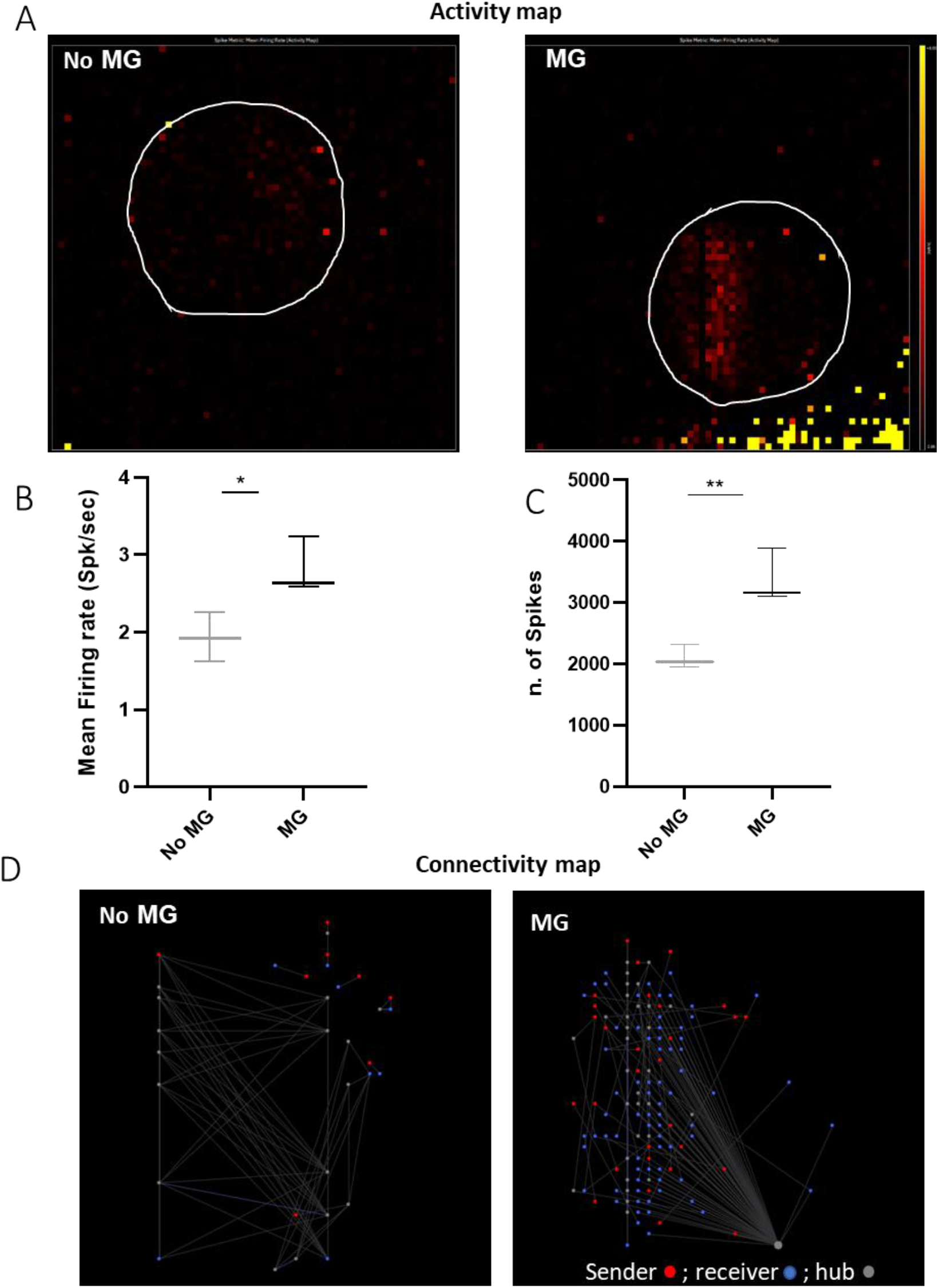
Neuronal functionality by HD-MEA in neuroimmune assembloids. A) Representation of the activity map in one hCO (cerebral organoid without microglia – noMG) and one neuroimmune assembloid (MG). The spots and red intensity represent spike activity recorded in the organoids. B-C) Quantification of the mean firing rate (number of spikes per second) and the number of spikes in hCOs (NoMG) and neuroimmune assembloids (MG). Results are expressed as a relative changes vs the reference electrode in the HD-MEA chip. D) Representation of the connectivity map in one hCO (cerebral organoid without microglia –noMG) and one neuroimmune assembloid. Red spots represent neurons sending electrical signals, blue spots represent neurons receiving electrical signals, and grey spots represent neuronal hubs involved in synchronization and burst propagation. Spot size represents the intensity of neuronal activity. Images are representative of three different experiments.

Microglial functionality in neuroimmune assembloids was evaluated following neuroinflammatory stimulation with 10 ng/mL lipopolysaccharide (LPS) for 24 hours. After 14 days of microglia assembly, transcriptional analysis of untreated and LPS-treated neuroimmune assembloids was evaluated by qRT-PCR (Fig. 7). Among the genes associated with neuroinflammation we found an evident modulation of CD45, IL1A, TNF α, NLRP3, CX3CR1, IL18, TMEM 119 and the post-synaptic marker PSD95 following LPS treatment. Surprisingly, all proinflammatory transcripts were upregulated following LPS treatment except for IL1A and IL18. LPS treatment induced a three-fold increase in CD45 expression and approximately two-fold increases in CX3CR1, NLRP3, TMEM119, and TNFA transcripts, whereas IL18 and IL1A transcript levels were reduced by the 50%.

**Figure 7.**
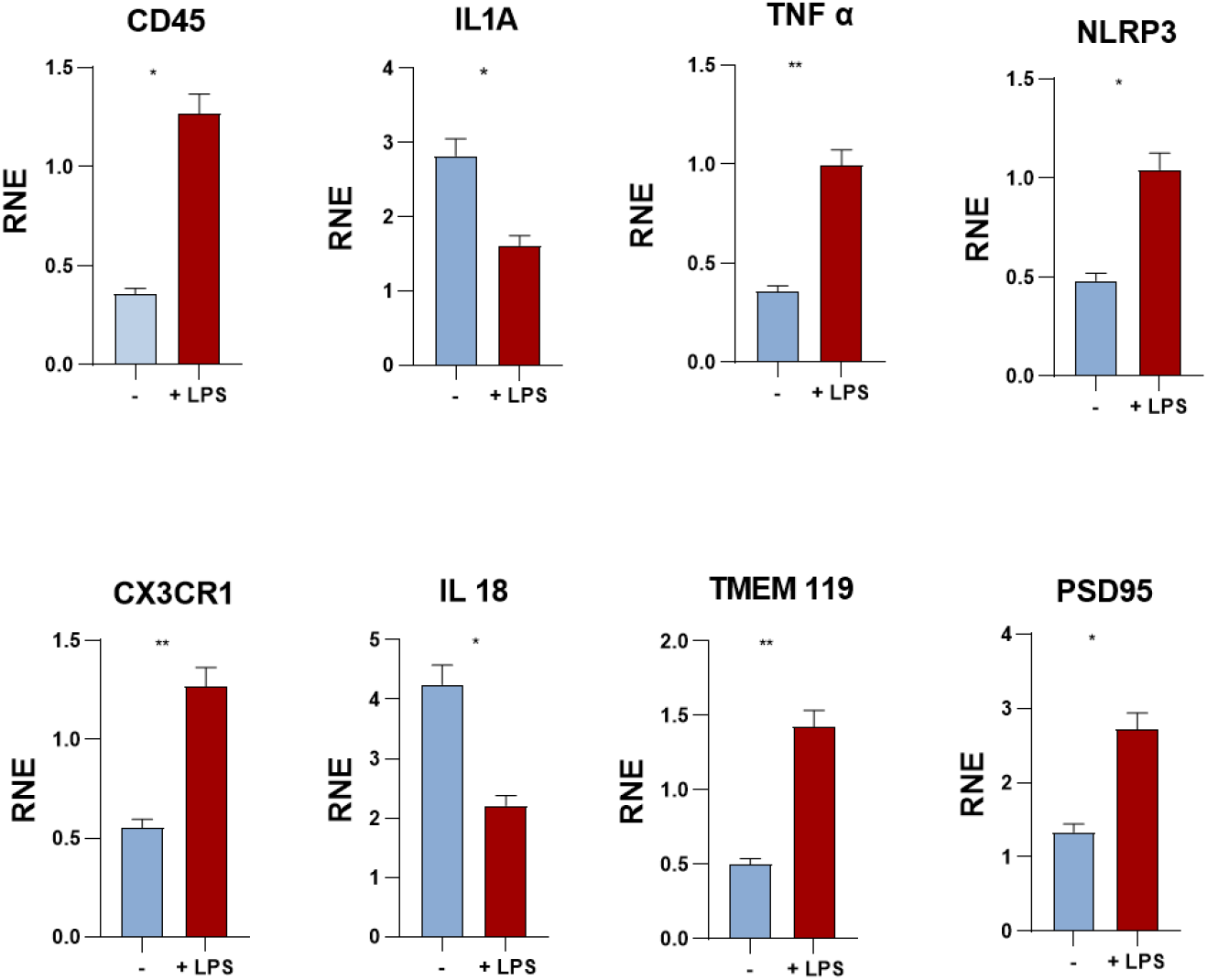
Gene expression of neuroimmune assembloids before and after LPS treatment. Semi-quantitative qRT-PCR analysis of cDNA isolated from neuroimmune assembloids after 14 days of microglia assembly (-) or after 24 hours of LPS treatment (+LPS). Relative transcript expression was calculated as ΔCt after normalization with GAPDH and represented as relative normalization expression (RNE). Data is representative of three different experiments.

Morphological analysis following Iba1 staining revealed that LPS-induced microglial activation promoted an amoeboid phenotype, as quantified by circularity analysis, with significant increases of 12.5% (Fig. 8A–B). After additional analysis we found that ramified microglia were more represented in control (no treated neuroimmune assembloids) whereas LPS treatment produced microglia cell loss in the external zone of the assembloid and amoeboids phenotype in the internal zones (data not shown).

**Figure 8.**
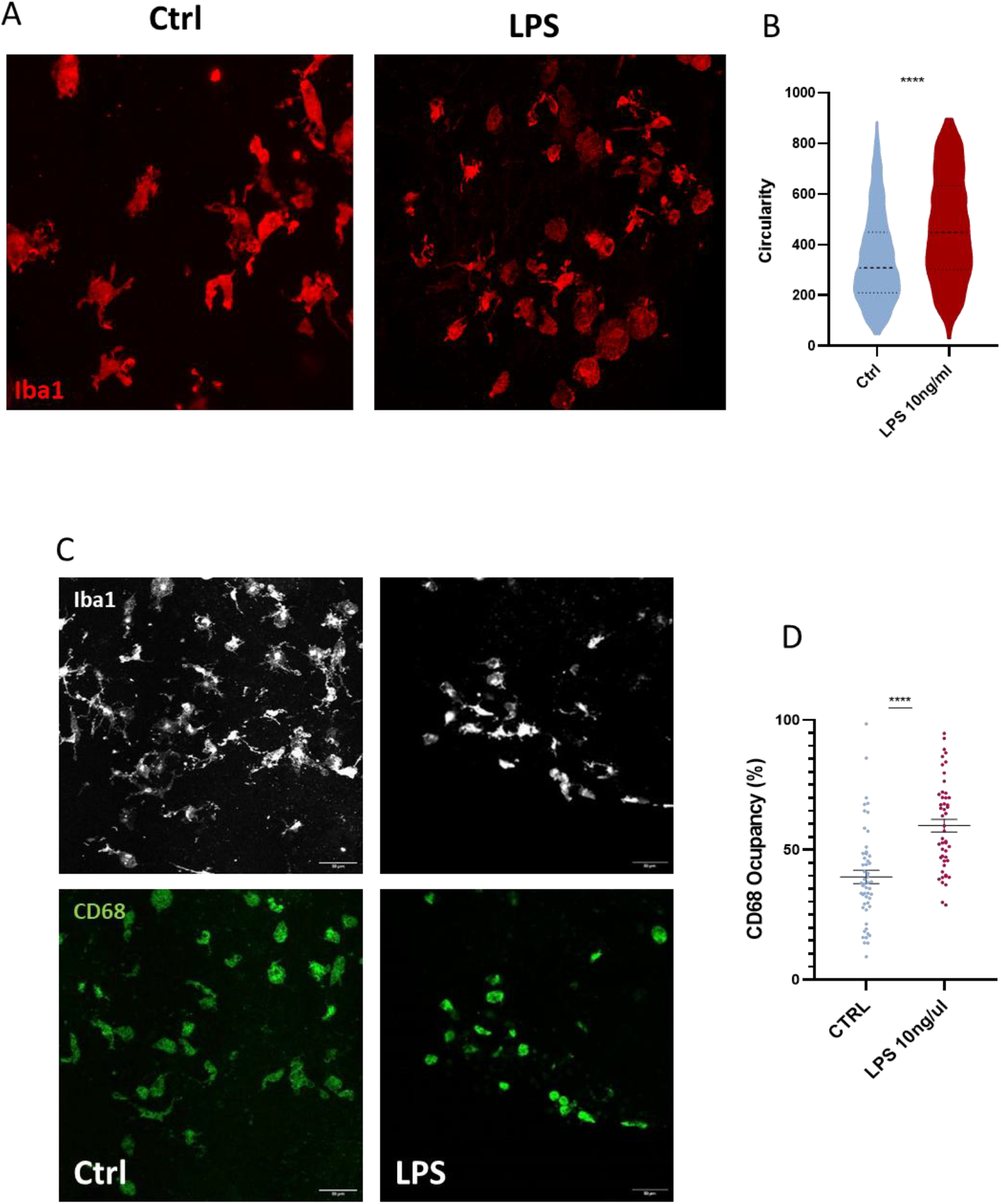
Microglia activation after LPS treatment by morphological changes and migration. A-B) Morphological analysis of microglia in neuroimmune assembloids 14 days after microglia assembly, before (Ctrl) and after 24 hours of 10ng/ml LPS treatment. Circularity was calculated in (B) after Iba1 staining using the specific plug-in of ImageJ software. C) Microglial activation by CD68 over-expression. Neuroimmune assembloids were stained with Iba1 (white) and CD68 (green) 14 days after microglia assembly and before (Ctrl) and after LPS treatment. D) CD68 occupancy was calculated in each Iba1-positive cell as the % of green staining (CD68) area relative to white staining (Iba1) area.

These LPS-induced morphological changes suggested an enhanced phagocytic state. CD68, a lysosomal protein upregulated in phagocytic microglia, was therefore used to further assess microglial activation. Immunostaining of neuroimmune assembloids with a CD68 antibody before and after LPS treatment showed a significant increase in CD68^+^ area occupancy within LPS-activated microglia, indicating upregulation of the phagocytic machinery in response to inflammatory stimulation (Fig. 8C-D). Phagocytic activity was further confirmed by 3D reconstruction in Video 1 that revealed a microglial cell engulfing an apoptotic nucleus.

## Discussion

The present study demonstrates the successful generation of neuroimmune assembloids through the incorporation of hematopoietic precursor cells into maturing brain organoids, resulting in functional microglia that migrate throughout the organoid structure and respond to inflammatory stimuli. This approach aligns with and extends recent advances in microglia-containing organoid models, while addressing several methodological considerations.

Brain organoids represent a powerful model for studying human brain development and genetic diseases, with promising applications in preclinical diagnostics, precision medicine [12], and AAV-based gene therapy [13]. However, despite their potential, brain organoids still exhibit methodological limitations that render them unsuitable for several areas of neuroscience research [14]. A notable limitation, particularly in guided brain organoids, where neural induction is driven by dual SMAD inhibition, is the absence of functional microglia. Unlike neurons, astrocytes, and oligodendrocytes, microglia originate from a distinct embryonic lineage. During late embryonic development, macrophage precursors migrate from the yolk sac into the brain parenchyma, where they proliferate, invade, and differentiate into mature, functional microglia. Microglia play a central role in brain development by shaping neuronal circuits, regulating synaptic transmission [15], and phagocytosing synapses, soluble antigens, cellular debris, and apoptotic cells. In pathological conditions and during brain aging, microglia also regulate neuroinflammation and astrocytic reactivity through the release of soluble inflammatory mediators, such as cytokines and chemokines. Thus, microglia are essential components for modeling neurodevelopmental processes, brain aging, and the pathophysiology of neurodegenerative diseases such as Parkinson’s and Alzheimer’s disease using brain organoids. Several models incorporating microglia into brain organoids have been proposed, yet they present significant limitations [8,14]. These include limited microglia lifespan [16], poor microglial colonization [17], insufficient or excessive phagocytic activity [3], and abnormal morphological reactivity [18]. Alternative approaches, such as co-culturing adult microglia with neural progenitor cells or reducing dual SMAD inhibition to promote endogenous microglial development, often result in increased heterogeneity and further complications.

Here, we present a neuroimmune assembloid model containing functional microglia derived from the same iPSC line used to generate the organoids. We define immunocompetence as the capacity of mature microglia to attain a functionally mature state, characterized by: (1) vigilance and complete migration with a mosaic-like distribution; (2) complex morphology responsive to inflammatory stimuli; (3) maintenance of homeostasis; and (4) effective phagocytosis coupled with a regulated inflammatory response. Our strategy to generate neuroimmune assembloids is based primarily on recapitulating *in vitro*, the developmental timing of microglial precursor migration. While current organoid cultures cannot fully replicate the chronological age of the developing brain, we employ early-stage brain organoids to include hematopoietic precursor cells, which can then differentiate *in situ* into mature microglia. After 30-40 days *in vitro*, our brain organoids continue to exhibit developmental characteristics, including the presence of neurogenic niches, expression of immature neuronal markers, and limited neuronal maturation. At this stage, mature astrocytes are only beginning to differentiate. Based on these observations, we infer that the organoids approximate the late fetal stage of brain development. To mimic hematopoietic precursor colonization and extend microglial lifespan within the organoids, we introduced undifferentiated microglial precursors following a brief *in vitro* priming period. Final microglial differentiation was then promoted by gradually transitioning the microglia from their specific culture medium to the brain organoid medium. Using this strategy, we were able to recapitulate key features of immunocompetent microglia within developing brain organoids. Microglia differentiated from iPSCs expressed multiple proteins associated with microglial homeostasis, such as PU.1 and P2Y12, among others. We also observed an increased release of specific cytokines into the culture medium, including IL-6, IL-1RA, IFN-γ and TNF-α (data not shown). These cytokines are known to be released at basal levels by astrocytes in the absence of microglia but are significantly upregulated in the presence of or by the microglia. Many protocols for incorporating microglia into brain organoids include the addition of CSF-1, TGF-β, and IL-34. These factors can be secreted by astrocytes during brain development and support microglial viability and maturation, along with other molecules such as cholesterol [19,20,21]. Bohlen et al. [19] demonstrated that astrocyte-conditioned medium could substitute for serum in microglial cell cultures, supporting viability while maintaining the microglia in a resting state without releasing inflammatory cytokines. For this reason, we decided not to include additional astrocyte-secreted factors, in order to reduce experimental variability arising from manipulation, batch differences, or supplier variations. Remarkably, within one week of assembly, these cells migrated to both the inner and peripheral regions of the organoids, thereby providing homeostatic support throughout the tissue. The morphological diversity observed, ranging from ramified to amoeboid forms, is consistent with microglial plasticity documented across multiple studies. Sabaté-Soler et al. [22] specifically noted both round and partially ramified microglial morphologies in their midbrain assembloids. Although we were unable to statistically quantify the reduction in the necrotic core, we did observe a visible decrease, along with evidence of microglial phagocytosis of apoptotic nuclei. More importantly, the differentiated microglia exhibited a clear response to a neuroinflammatory stimulus induced by LPS treatment. Specifically, we observed a pronounced morphological shift toward a rounded, amoeboid form, typical of activated, migrating microglia with phagocytic potential accompanied by the modulation of several proinflammatory transcripts. Surprisingly, we observed an LPS-induced upregulation of all pro-inflammatory transcripts tested except for IL1A and IL18. This would suggest a differential time-scale regulation of IL1A and IL18 during the early phases of the inflammatory response and coincides with the early death of microglia in the peripheral region of the assembloid (Fig. 5D). An increase in phagocytic activity was further supported by elevated expression of CD68, a lysosomal transmembrane protein involved in the uptake and degradation of cellular debris as similarly reported by both Lecca et al. [23] and Xu et al. [3], validating this as a reliable marker of microglial activation in 3D organoid systems. Importantly, microglia inclusion enhanced neuronal network complexity as evidenced by electrophysiological analyses, implying their crucial role in synaptic maturation and circuit functionality, consistent with emerging literature highlighting microglia’s influence on neuronal development and network refinement [22,24].

Integration of iPSC-derived microglia into these organoids to form neuroimmune assembloids represents a significant advance in modeling brain development and neuroimmune interactions, addressing the critical absence of immune cells in traditional organoids. Our results confirm the functional fidelity of iPSC-derived microglia in this 3D context, providing a physiologically relevant model for neuroinflammation studies and neuroimmune crosstalk in human brain tissue. Even with some limitations, such as a relatively short functional lifespan, sparse myelination despite the presence of oligodendrocyte lineage markers, and the lack of blood-brain barrier cell types, this platform offers a promising tool for investigating neuroimmune mechanisms underlying brain development, homeostasis, and neuropathologies. The slight decrease in total microglial area at 14-30 days compared to 7 days may reflect several non-mutually exclusive mechanisms. It could indicate phagocytic clearance of excess microglia by surviving cells, a homeostatic mechanism observed *in vivo*, or it may represent death of microglia that failed to establish productive interactions with neural cells, a selection process that would enrich for functional populations.

Overall, the combination of well-characterized cerebral organoids and immunocompetent microglia within them advances the field by recapitulating both neuronal and immune components of the brain, enabling the study of cellular interactions during development and disease modeling with high translational potential.

## Supporting information

Phagocytosis video 1

## Acknowledgment

This work was supported and funded by the Spanish Ministry of Science, Innovation and Universities “Programa de Generación de Conocimiento” (PID2023-146826OB-I00, PID2022-140236OB-I00, CNS2025-166813). P.R.G. was funded by an IKUR2030 program from the Basque Government. N.U.A. holds a Basque Government predoctoral fellowship, and I.M.F. holds a contract from the Dep. of Industry of Basque Governement under the programme ELKARTEK (KK-2025/00089).

The authors acknowledge the support of the ACHIEVE Platform at Achucarro Basque Center for Neuroscience (*Cell Analytics/Biomarkers/Molecular Biology/Human Biomodels/In Vivo Phenotyping Unit/Imaging*) for technical assistance and access to advanced infrastructure. SHSY5Y VAMPIRE cells for phagocytosis assay were kindly provided by Dr. A. Sierra (Achucarro Basque Center for Neuroscience). Protocol schemes in Figure 1-3 were generated using Figurelabs, an online platform for scientific figure creation.

## Figure legends

**Video 1**. Z-Stack caught with a 63X objective and 3x zoomed with 3D reconstruction of two microglia cells in a neuroimmune assembloid, 14 days after assembly. Nuclei are counterstained with DAPI. The video shows how microglia cells stained with Iba1 (red) and CD68 (green) are phagocyting an apoptotic condensed blue nucleus.

